# The FusX TALE Base Editor (FusXTBE) for rapid mitochondrial DNA programming of human cells *in vitro* and zebrafish disease models *in vivo*

**DOI:** 10.1101/2021.05.18.444740

**Authors:** Ankit Sabharwal, Bibekananda Kar, Santiago Restrepo-Castillo, Shannon R. Holmberg, Benjamin Luke Kendall, Ryan P. Cotter, Zachary WareJoncas, Karl J. Clark, Stephen C. Ekker

**Author notes:** **Corresponding author:** Stephen C. Ekker-, Department of Biochemistry and Molecular Biology, Mayo Clinic, 200 1^st^ Street SW, Rochester, MN 55905. **E-mail IDs: AS**-; **BK**-; **SRC**-; **SRH**-; **BLK**-; **RPC**-; **ZWJ**-; **KJC**-; **SCE**.

## Abstract

Functional analyses of mitochondria have been hampered by few effective approaches to manipulate mtDNA and a lack of existing animal models. Recently a TALE-derived base editor was shown to induce C-to-T (or G-to-A) sequence changes in mtDNA. We report here the FusX TALE Base Editor (FusXTBE) to facilitate broad-based access to TALE mitochondrial base editing technology. TALE Writer is a *de novo in silico* design tool to map potential mtDNA base editing sites. FusXTBE was demonstrated to function with comparable activity to the initial base editor in human cells *in vitro*. Zebrafish embryos were used as a pioneering *in vivo* test system, with FusXTBE inducing 90+% editing efficiency in mtDNA loci, the first example of majority mtDNA heteroplasmy induction in any system. Gene editing specificity as precise as a single nucleotide was observed *in vivo* for a protein-coding gene. Non-destructive genotyping enables single animal mtDNA analyses for downstream biological functional genomics applications. FusXTBE is a new gene editing toolkit for exploring important questions in mitochondrial biology and genetics.

## Introduction

Mitochondria have critical roles in cellular homeostasis, metabolism, apoptosis, and aging^1^. Approximately, 80% of the adult-onset and 25% of the pediatric-onset mitochondrial disorders can be attributed to genetic lesions in the mitochondrial genome. Mutations in both nuclear DNA as well as mitochondrial DNA (mtDNA) are pathogenic^2,3^. For example, a recurring 5 kilobase deletion underlies Kearns-Sayre syndrome (KSS), and a single point mutation in mtDNA is the basis for mitochondrial encephalopathy, lactic acidosis, and stroke-like episodes (MELAS) syndrome^4,5^. The complex genetics, including penetrance and phenotypic heterogeneity observed in the case of mtDNA-associated disorders, can be explained by the homoplasmic or heteroplasmic condition of the cell. A prototypical cell harbors around 100-100,000 copies of circular, double-stranded mtDNA molecules. The proportion of the mutant to wild type mtDNA within a cell governs the progression of the mitochondrial disease, the severity of which is directly proportional to the number of mutant mtDNA molecules. Despite this clear association of genotype with the disease, there are no current treatments for patients with mitochondrial disease. Understanding the pathophysiology of mitochondrial disorders contributed by variations in the mtDNA has been impeded due to the lack of an effective toolkit to engineer the powerhouse of the cell. The lack of systems for the quantitative delivery of exogenous DNA or RNA to mitochondria also restricts approaches for mtDNA editing. While gene-editing tools such as CRISPR and TALENs enable the rapid manipulation of the nuclear genome^6–10^, there are few tools to manipulate/engineer vertebrate mitochondrial DNA (mtDNA) mainly due to the absence of any common DNA repair mechanisms. DNA nucleases that create double-stranded breaks in the nuclear genome and subsequently recruiting repair machinery, instead are proposed to induce the degradation of mtDNA^11–14^. These inherent mitochondrial gene editing challenges have been circumvented by expanding on the existing protein-based gene editing delivery systems. Currently, few strategies take advantage of this feature to target and degrade pathogenic mtDNA via double-strand breaks.

Gene editing tools, such as mitochondrially targeted zinc-finger nucleases (mtZFNs), and mitochondrially targeted transcription activator-like effector nucleases (mitoTALENs) have been reported to shift mtDNA heteroplasmy in models of mitochondrial disorders^11, 12, 15^. Mitochondrially targeted nucleases selectively bind the mtDNA harboring deletions/edits and introduce double stranded breaks, thereby triggering rapid degradation of mtDNA. In a previous study by our group, we developed an alternative approach to introduce precise deletions in the mtDNA using a novel “block and nick” approach to model KSS in zebrafish^16^. However, the utility of these nuclease-based gene editing systems cannot be extended to correct or model homoplasmic and single nucleotide pathogenic mtDNA variations. Recently, Mok *et al*. described a novel base editing system in the mitochondria^17^ using a bacterial cytidine deaminase toxin, DddAtox. The present system uses split halves of DddAtox fused to each of the TALE monomers programmed to bind a specific sequence on mtDNA. Mitochondrial DdCBE (DddA-derived cytosine base editor) introduces C-to-T base edits in the spacer region between the two TALE arms with optimal editing efficiency *in vitro*. This system has a strong sequential preference for the presence of T at the 5’ end of the C around the targeted editing site. Further, Mok *et al*. show that to target mtDNA DddAtox can be split into two halves at two different positions, each of the split-halve molecules swapped with any one of the TALE monomers. Using this as a critical study design constraint, one needs to test four different combinations of the DdCBE to interrogate a single target, a process that can be quite time-consuming if extended to edit multiple targets present on a given mtDNA molecule. The application of this editing system has been extended to *in vivo* in a study wherein Lee *et al*., have demonstrated to introduce base edits in the mice mtDNA utilizing DddA-TALE fusion deaminases. C-to-T transition was detected in multiple tissues and was also observed to be germline transmitted in two F1 offsprings born to edited F0 animal^18^.

Here we describe a universal and programmable vertebrate mtDNA engineering toolbox suitable for the establishment of new *in vitro* cell and *in vivo* animal models. We build upon the existing DdCBE system to develop a rapid, accessible, and efficient programmable FusX^19^ TALE base editor system with high editing efficiency *in vitro* and *in vivo*. To predict all potential editable sites in mtDNA, we have designed a customizable script that also determines the target sequences amenable to premature termination codon (PTC) induction via C-to-T base editing. A non-invasive method well adapted to zebrafish provides a unique advantage to genotype and propagate injected embryos that are harbor up to 90% edits in their mtDNA. The findings from this study provide insights in exploring the mitochondrial biology and genetics of disorders caused by mutations in mtDNA.

## Materials and Methods

### Generation of mitoFusX TALE Base Editor (FusXTBE)

For the generation of this next generation human mitochondrial TALE base editor, the cassette containing the N-terminus half of the DddA_tox_ (split at 1397 amino acid position), uracil glycosylase inhibitor (UGI) and 3’UTR of human *ATP5B* was amplified from the published DdCBE construct (Addgene plasmid no. 157843). The PCR product was digested with the restriction enzymes XbaI and EcoRI. Digested DNA fragments were then subsequently cloned in the pkTol2c-*COX8A*-mTALEN vector linearized with XbaI and EcoRI to obtain pkTol2c-**FusXTBE-N**. To generate pkTol2c-FusXTBE-C, the cassette containing the C terminus half of the DddA_tox_ (split at 1397 amino acid position) and UGI was amplified from the DdCBE plasmid (Addgene plasmid no. 157844). The PCR amplicon was digested with XbaI and BsU36I restriction enzyme and cloned in the pkTol2c-FusXTBE-N vector linearized with XbaI and Bsu36I to generate pkTol2c-**FusXTBE-C** plasmid. Clones were confirmed by Sanger sequencing (GENEWIZ LLC, USA). For zebrafish mitochondrial FusX TALE base editor constructs, individual cassettes including partial N- and C-terminus of TALE domain, along with DddA_tox_ (split at 1397 amino acid position), and UGI were ordered as clonal fragments in pTwist Amp High Copy plasmids from Twist Bioscience, USA. Each of the clonal fragments was then separately cloned in the BclI-PstI cut pk*idh2*GoldyTAL-lacZ-Fok1 vector backbone, generating plasmids **pT3-FusXTBE-N and pT3-FusXTBE-C**. The final FusX mitoTALE-DddA_tox_ backbone plasmids will be made available via our Addgene repository. RVDs specific to prioritized mtDNA genetic loci (Supplementary table 1) were then cloned in the receiving vector backbones using the one-step FusX assembly system^19^ (Figure 1A). Each of the constructs was sequence-verified by Sanger sequencing. FusX Extender series was build using previously published FusX system where an additional RVD in all possible combinations was added to each of the pLR (last half-repeat) plasmids. This modified design enables us to synthesize TALEs binding to sequences 17 bp long on mitochondrial or nuclear genome.

**Figure 1:**
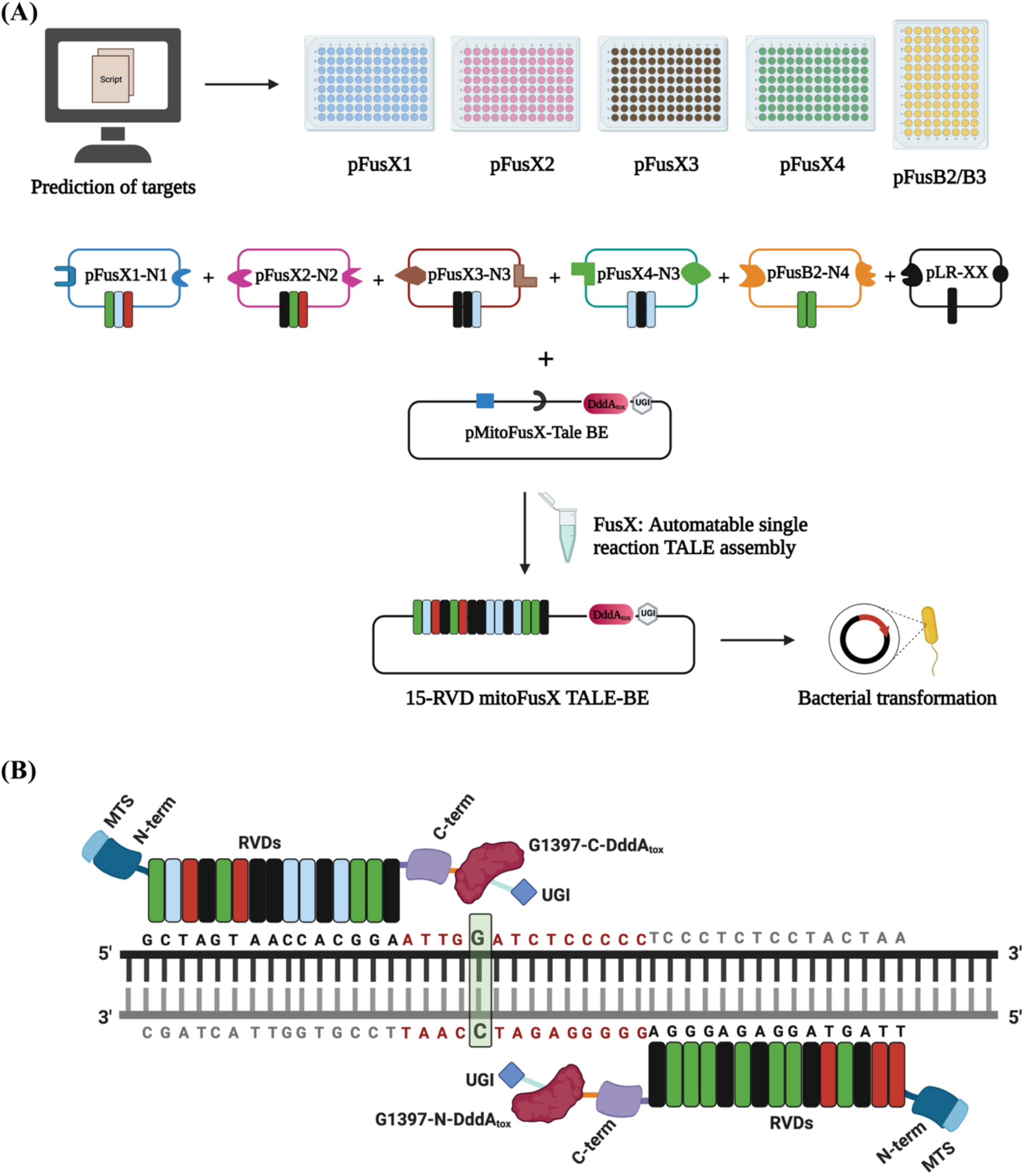
Schematic for the assembly of mitochondrial TALE base editors by the FusX reaction with the FusXTBE architecture. **(A)** FusX library of all combinations of RVD dimers and trimers enables the mitochondrial TALE base editor to be built in a single Golden Gate cloning step. **(B)** Schematic of FusXTBE binding to the human mitochondrial DNA target, ND4. Repeat divariable residues are shown in different colors (Green-Guanine-NN; Blue-Cytosine-HD; Red-Thymine-NG; Black-Adenine-NI). DNA binding region for each of the TALE monomers is black in color, and the spacer region is depicted in red. Genomic target C is highlighted in green with bold and italicized font. Figure 1 was created using Biorender.com

### TALE-based technology design tool for mitochondrial genome engineering

TALE Writer, a script for the efficient design of TALE-based technologies, was developed using Python 3.7.3. The script can be used to (I) design TALEs around user-specified target sequences, (II) design C-to-T base editors against 5’-T**C**-3’ loci^17^, and (III) design C-to-T base editors specifically for premature termination codon (PTC) induction^20^. The main input of the script is a sequence of target genes, and the output is the top-ranked design permutation (for TALE design) or a list of the top-ranked design permutations for each target within the input sequence (for C-to-T base editor design). In general, a user-specified sequence is analyzed considering three design parameters: (I) Obligatory 5’-T^0^ for all TALE repeat arrays^19^, (II) desired cut site or edit site at least four base pairs from the target sequence of either TALE repeat array, and (III) target and spacer sequences between 14 and 16 base pairs long (these lengths can also be specified by the user). Since there can be multiple design permutations for every target, in order to decrease the size of the output, a scoring system was developed; each possible design permutation is given a score, consisting of the sum of four Gaussian functions that depend on the length of the target and spacer sequences, and the position of the desired cut site or edit site within the spacer region. In brief, permutations that follow a 15-15-15 design format (TALE-spacer-TALE) and that possess the desired cut site or edit site in the middle of the spacer sequence are favored. For PTC induction, herein defined as a DNA edit resulting in the generation of a non-functional gene product due to the premature termination of translation^20^, three types of edits were identified. First, the in-frame 5’-TGA-3’ motif, in which a G-to-A edit (equivalent to a C-to-T edit on the opposite strand) results in the stop codon UAA. Second, the 5’-TCAA-3’ motif (CAA in-frame), in which a C-to-T edit also results in the stop codon UAA. Lastly, the 5’-TCAG-3’ motif (CAG in-frame), in which a C-to-T edit results in the stop codon UAG. Noteworthy, PTC induction should be possible both in nuclear DNA and mitochondrial DNA through any of these edits. It is also worth mentioning that the availability of types of edits is currently limited by the strong 5’-TC-3’ context preference of the first generation of DdCBEs^17^. TALE Writer is hosted at https://github.com/srcastillo/TALE-Writer under a GNU General Public License v3.0.

### Transfection of the mammalian cell line with mitoFusX TALE Base Editor

HEK293T cells were cultured and maintained at 37°C with 5% CO2. HEK293T cell line was obtained from ATCC and cultured in DMEM GlutaMax media (Thermo Scientific, USA) supplemented with 10% fetal bovine serum (FBS) (Gibco, USA) and 1X Penicillin-Streptomycin solution (Pen-Strep) (Thermo Scientific, USA). 30,000 cells/well were seeded in a six-well plate 18 hours prior to the time of lipofection by Lipofectamine3000 (Thermo Scientific, USA). Cells were transfected with 750 ng of each TALE-BE monomer (left arm-*MT-ND4* pkTol2c-**FusXTBE-C** and right arm-*MT-ND4* pkTol2c-**FusXTBE-N**) and 500 ng of pkTol2C-EGFP as a positive control for the lipofection. Media was changed 24 hours post-transfection and cells were cultured for 72 hours before collection for mitochondrial genotyping.

### Genomic DNA isolation and mitochondrial genotyping

Following 3 days post-transfection, media was aspirated, and cells were washed with 1X phosphate-buffered saline (PBS) (Thermo Scientific, USA). Total genomic DNA was extracted from cells using DNeasy Blood and Tissue Kit (Qiagen, USA). Primers flanking the *MT-ND4* gene were used to amplify the edited locus using MyTaq polymerase (Bioline, USA). PCR amplicons were gel extracted and purified using QIAquick Gel Extraction Kit (Qiagen, USA). Purified samples were submitted to Genewiz (GENEWIZ LLC, USA) for Sanger sequencing. Mitochondrial heteroplasmy was quantified using EditR software^21^ to predict the percentage of edits in the m.G11922 site in the *MT-ND4* gene. Raw Sanger sequencing files in .ab1 format were uploaded for each of the control and non-transfected samples.

### Zebrafish handling and husbandry

Adult zebrafish and embryos were maintained according to the guidelines established by the Mayo Clinic institutional animal care and use committee (IACUC number: A34513-13-R16).

### Zebrafish microinjection

Targets predicted by TALE Writer were prioritized to introduce edits in protein-coding and tRNA genes. Mitochondrially encoded cytochrome c oxidase I (*mt-co1*) (non-synonymous substitution), mitochondrially encoded cytochrome c oxidase III (*mt-co3*) (pre-mature termination induction), and mitochondrially encoded tRNA-leu (*mt-trnl1*) were chosen as target sites in the zebrafish mtDNA locus. *mt-co1* and *mt-co3* code for subunits of complex IV of the mitochondrial respiratory chain, whereas *mt-trnl1* is involved in tRNA aminoacylation. Sequence-verified mitoTALE-BE clones were linearized with PstI restriction enzyme. Linearized plasmid was used as a template to *in vitro* transcribe mRNA using T3 mMessage mMachine transcription kit (Thermo Scientific, USA). mRNA was purified using NEB Monarch RNA cleanup kit (NEB, USA). 3 nl of 40 pg mRNA of each arm of the FusX mitoTALE-BE (left-**pT3-FusXTBE-N** and right-**pT3-FusXTBE-C**) was injected into single-cell zebrafish embryos. Injected embryos were then raised in embryo water at 29°C.

### Non-invasive enzymatic zebrafish genotyping

To assess the *in vivo* editing efficiency of the mitoTALE BE, a method of genotyping was needed that enabled the rapid investigation of the functional readout of mtDNA edits. A non-invasive enzymatic method to extract nuclear DNA from zebrafish embryos was employed with modifications^22^. Larvae aged 3 days post fertilization (dpf) were rinsed thrice in fresh embryo media, followed by three washes in collection buffer (30 mM Tris-HCl, 1X tricaine solution). Larvae were aspirated in 40 μL of genotyping buffer (1.5μg/mL proteinase K, 30 mM Tris-HCl, 1X tricaine solution) into a 96 well plate and were incubated in a shaker at 37°C for 20 minutes. After incubation, 15μL of the lysis buffer (10 mM Tris-HCl, pH 8.0, 50 mM KCl, 0.3% Tween-20, 0.3% NP-40 and 1 mM EDTA) was added to the incubation mix. The solution was incubated at 98°C for 5 minutes to deactivate the proteinase K and release DNA from cells shed by the larvae upon agitation. This DNA was used as a template for the PCR amplification to genotype the larvae. Each larva was then individually rescued from the 96 well plate and rinsed in 4-quadrant petri dishes with fresh embryo water, before being placed in 24-well plates for incubation at 29°C.

### Mitochondrial DNA genotyping in zebrafish embryos

PCR amplification was performed for each of *mt-co1, mt-co3, mt-trnl1* using the respective primers mentioned in supplementary table 2. PCR reaction consisted of 4 μL of 5x MyTaq Red PCR Buffer, 1 μL of 10μM forward primer, 1 μL of 10μM reverse primer, 0.2 μL of MyTaq DNA Polymerase, 8 μL of template, and 5.8 μL dH2O. Samples were run in a thermal cycler with the following PCR conditions: (1) 95°C, 3 mins; (2) 95°C, 1 min; (3) 58°C, 30 sec; (4) 72°C, 30 sec; (5) go to step 2 34X; (6) 72°C, 5 min; (7) 12°C, hold. PCR amplicons were run on 2% agarose gel and analyzed. 8 μL of the PCR amplicon was used as the substrate DNA for the restriction fragment length polymorphism (RFLP) assay in a total reaction volume of 20 μL (1X buffer, 5 units of restriction enzyme, and remaining volume dH2O). Embryos injected with *mt-col, mt-co3*, and *mt-trnl1* TALE BE RNAs were surveyed for loss of BstYI, gain of MseI, and loss of SacI restriction sites, respectively. PCR amplicons for the samples confirmed positive by RFLP were then gel extracted and purified using the QIAquick Gel Extraction kit (Qiagen, USA). Purified amplicons were then submitted to Sanger sequencing (GENEWIZ LLC, USA) to estimate the editing efficiency and any off-target edits in the protospacer region (protospacer region is the sequence between two bound TALE monomers). Chromatogram files were then analyzed using EditR software^21^ to calculate the heteroplasmy in the injected embryos. Embryos harboring base edits were then placed on the fish facility at 6 dpf to enable the investigation of potential germline transmission and tissue-specific heteroplasmy of the edits.

## Results

### Assembly of FusXTBE plasmids

To assess the activity of FusXTBE plasmids in mammalian cells, we cloned the split halves of DddA_tox_ (split at 1397 amino acid position) and UGI in a FusX-compatible, codon-optimized destination plasmid, pkTol2c-*COX8A*-mTALEN. This receiving plasmid contained the mitochondrial targeting sequence of human cytochrome c oxidase subunit 8a (*COX8A*) upstream of the N-terminal domain of the TALE cassette to facilitate the import of base editor inside mitochondria. The resulting MTS-TALE-DddA_tox_ module was cloned upstream to 3’UTR sequence from the human *ATP5B* gene in order to increase the efficiency of mRNA sorting near the proximity of mammalian mitochondria^23^. To quantify the editing efficiency in vertebrate zebrafish embryos, a cassette containing DddA_tox_ was cloned at the 3’ end of the C-terminus of the TALE domain in the linearized pk-*idh2*GoldyTAL-lacZ-Fok1 destination vector. We identified a highly functional and sufficient mitochondrial targeting sequence from a reverse engineered *in vivo* protein trap in a nuclearly encoded mitochondrial protein^16,24^. This derived minimal mitochondrial targeting sequence (MTS) was highly active in zebrafish *in vivo* compared to previously described MTS sequences and was used to target an array of different proteins capable of interacting with the mtDNA genome^16^. We were able to rapidly assemble and test different TALE constructs using the previously published FusX system^19^. This method utilizes a library of dimer and trimer RVD (TALE repeat) modules that uses Golden Gate cloning to synthesize a TALE that targets any 15- and 16-mer DNA sequence of interest (Figure 1A-B).

### TALE Writer predicts potential base editing sites in the human and zebrafish mitochondrial genome

TALE Writer, a computational tool for the efficient design of TALE-based technologies, was developed utilizing Python 3.7.3 to rapidly identify potential target sites for base editing in the human and zebrafish mitochondrial genomes. Based on specific design parameters from prior TALE-based nuclease and gene editor work (described in the methods section), the script allows the identification of cytosines within 5’-T**C**-3’ loci that can potentially be targeted by FusXTBEs. Some of these edits can lead to the induction of premature termination codons (PTCs), which can result in the generation of non-functional gene products due to the premature termination of translation^20^. TALE Writer can specifically identify such targets and calculate their possible FusXTBE design permutations contingent on user-specified parameters, such as the lengths of the TALE arms and the spacer region. Moreover, to decrease the output size, a scoring system that favored a 15-15-15 design format (TALE-spacer-TALE) with target cytosines within the middle of the spacer region was adopted.

Following user-specified design parameters, TALE Writer identified a total of 1,547 potential target sites for C-to-T base editing via FusXTBEs in the human mitochondrial genome, 89 of which were amenable to PTC induction (Figure 2A). For example, multiple design permutations were found for the locus m.G11922, which coincides with an in-frame 5’-TGA-3’ motif in the forward strand of the human gene *MT-ND4*. In this case, a successful G-to-A edit (equivalent to a C-to-T edit on the opposite strand) leads to the stop codon UAA. However, this target site is only 215 bp upstream of the natural termination codon of the 1,378 bp-long *MT-ND4* gene (Supplementary figure 1A), which could complicate downstream analyses of base editing.

**Figure 2:**
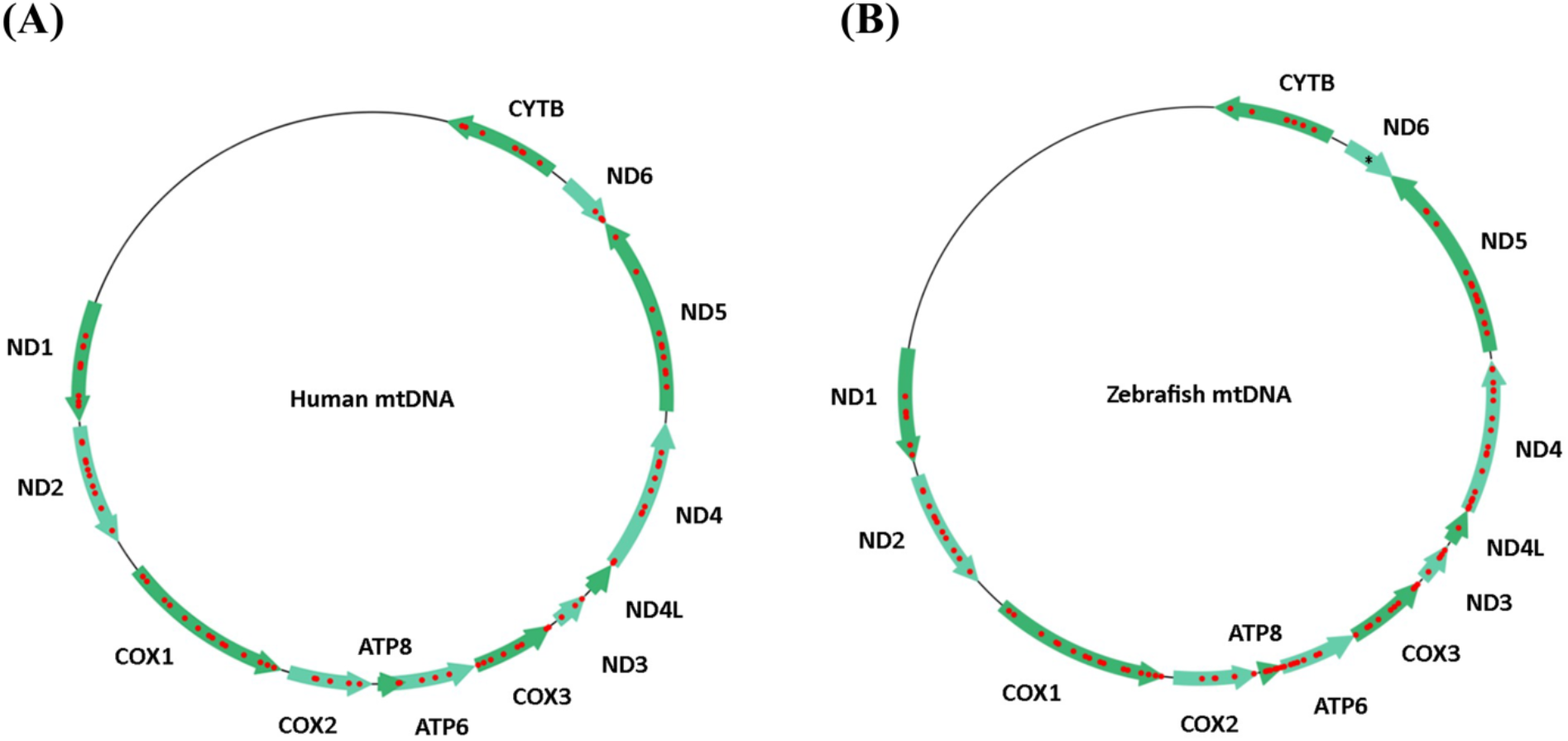
Predicted targetable sites for premature termination codon (PTC) induction in human and zebrafish mtDNA as determined by TALE Writer. **(A)** Using design parameters extrapolated from TALEN and initial mtDNA base editor work and including assembly parameters for the FusX system, the human mitochondrial genome possesses a possible 1,547 target sites for C-to-T base editing, 89 of which are amenable to PTC induction (red circles). **(B)** Similarly, the zebrafish mitochondrial genome possesses a total of 1,513 target sites for C-to-T base editing, 112 of which correspond to targets for PTC induction (red circles). The single black asterisk in the zebrafish gene mt-nd6 represents a site for PTC induction made available by setting the maximum TALE arm and spacer lengths to 17 bp; such a FusXTBE format can be assembled via our FusX extender kit-as a single-tube reaction.

Similarly, a total of 1,513 potential target sites for C-to-T base editing via FusXTBEs were found in the zebrafish mitochondrial genome, 112 of which coincided with loci amenable to PTC induction (Figure 2B). For example, multiple design permutations were found for the locus m.C10215, which coincided with a 5’-TCAA-3’ motif in the forward strand of the zebrafish gene *mt-co3*. In this case, a successful C-to-T edit leads to the stop codon UAA. Although the top-ranked design permutation for this target had a low-score according to our adopted scoring system (Supplementary figure 1B). This target site is located 305 bp upstream of the natural termination codon of the 786-bp long *mt-co3* gene, which could facilitate downstream analyses of base editing.

Based on the originally selected design parameters, no targetable sites amenable to PTC induction in the zebrafish gene *mt-nd6* were found (Figure 2B). However, by expanding the allowable lengths of TALE arms and spacer regions up to 17 bp, a 16-16-17 (TALE-spacer-TALE) design permutation for PTC induction was identified in this gene (Figure 2B). Specifically, the zebrafish locus m.C14967, which coincides with an in-frame 5’-TGA-3’ motif in the heavy strand of the gene *mt-nd6*, can be targeted. A successful G-to-A edit in this site (equivalent to a C-to-T edit in the opposite strand) would result in the UAA stop codon. Our FusX Extender series system can enable the manufacturing of such a design permutation as a single-tube reaction.

### Engineering base edits *in vitro* using FusXTBE

To assess the activity of the modular mito FusXTBE system *in vitro*, we used the computational tool TALE Writer, to predict the target sites in the human mtDNA locus that are amenable to PTC induction. Out of the 89 potential predicted target sites, we chose to edit the cytosine on the reverse strand at the m.G11922 position of *MT-ND4* gene (Figure 3A). We transfected HEK293T cells with each of the monomers of the FusX compatible TALE base editor with G1397-C-DddA_tox_ (left) and G1397-N-DddA_tox_ (right) and measured the editing efficiencies at 3 days post-transfection. PCR across the target locus and subsequent Sanger sequencing revealed editing efficiency at C_5_ in the range of 30 ± 4% (Figure 3B-C). No non-specific edits were observed in the spacer region between the two TALE binding sites, despite favorable C_1_ and C_8_ present upstream and downstream to the PTC target. To gain a better understanding of the effect of the programmable FusX-TALE modular domain on the editing activity, we compared the FusXTBE system with the original DdCBE construct reported^17^ targeting the same mtDNA locus. Cells transfected with the original DdCBE system exhibited editing efficiencies in the range of 35 ± 3 % across a series of experiments (Figure 3B-C). These results indicate that the FusXTBE system has a comparable editing efficiency *in vitro* when bound to the mitochondrial DNA and established this tool for potential additional applications, leveraging the informatic base mapping and rapid assembly system.

**Figure 3:**
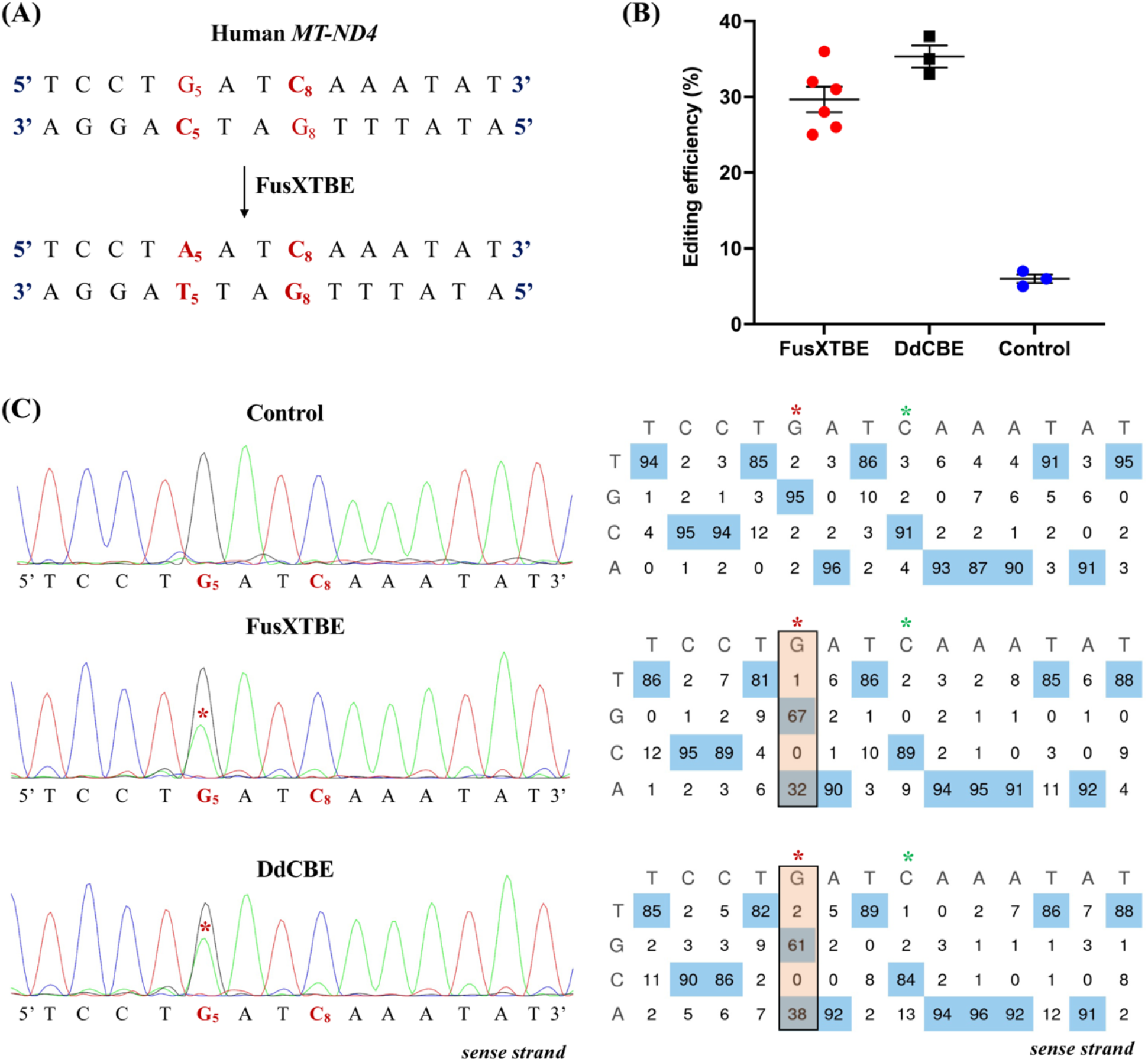
Engineering base edits in human cells *in vitro* using the FusXTBE system. **(A)** Potential target sites favorable for base editing in the protospacer region of the targeted human *MT-ND4* locus. **(B)** Editing efficiency assessed of mitochondrial FusXTBE and DdCBE system in HEK293T cells, assessed by Sanger sequencing. Data is represented from independent experiments. Black circle represents editing efficiency by FusXTBE system and black circle denotes DdCBE system. Blue circle represents the wild type heteroplasmy in the target locus in the non-transfected cells. Error bars are represented as standard error of the mean. No significant difference was observed in the editing efficiencies between the FusXTBE and DdCBE system (p-value >0.05 as determined by Student’s t-test). **(C)** Representative chromatogram of the control (non-transfected) and cells transfected with FusXTBE and DdCBE plasmids. Asterisk (*) denotes the site of edit (C_5_ position) with corresponding editing percentage (C-to-T or G-to-A). Chromatograms and editing table plot were obtained using EditR.

### FusXTBE introduces base edits *in vivo*

We used TALE Writer to predict potential sites amenable to base editing in the protein-coding and tRNA genes in zebrafish mtDNA. Out of the 1,513 potential sites, we prioritized three targets: two protein-coding genes, *mt-co1* and *mt-co3*, and the *mt-tRNA-leu* gene (or *mt-trnl1*). Amino acid sequences of the zebrafish Mt-Co1 (Figure 4A) and Mt-Co3 (Figure 4B) proteins displayed more than 80% similarity to their human orthologs indicating a high level of sequence conservation between these vertebrate species. Zebrafish *mt-trnl1* displayed ~69% sequence similarity with the human ortholog at the nucleotide level (Figure 4C). The zebrafish-applicable version of the FusXTBE specific to each of the target loci were assembled using the FusX assembly protocol. The expression of the mitochondrially targeted TALE base editor system (left-pT3-**FusXTBE-N** and right-pT3-**FusXTBE-C**) as *in vitro* synthesized mRNA was injected into the single-cell zebrafish embryos (Figure 4D). More than 70% survival was observed for the embryos injected with *mt-co1, mt-co3* and *mt-trnl1* BE RNAs over a range of experiments. 3 dpf injected embryos were subjected to non-invasive genotyping to facilitate downstream, longitudinal analyses during the future course of studies (Figure 4D). Survival was monitored till 2 days post genotyping and mortality in the range between 15-20% was observed across the three target-specific genotyping experiments. Subsequently, DNA was extracted from the shed cells in the solution and was used as a template for downstream assays such as PCR and RFLP. Base edits in any of the target sites either created a loss or gain of restriction enzyme site, facilitating the scoring for embryos harboring variations in the mtDNA locus.

**Figure 4:**
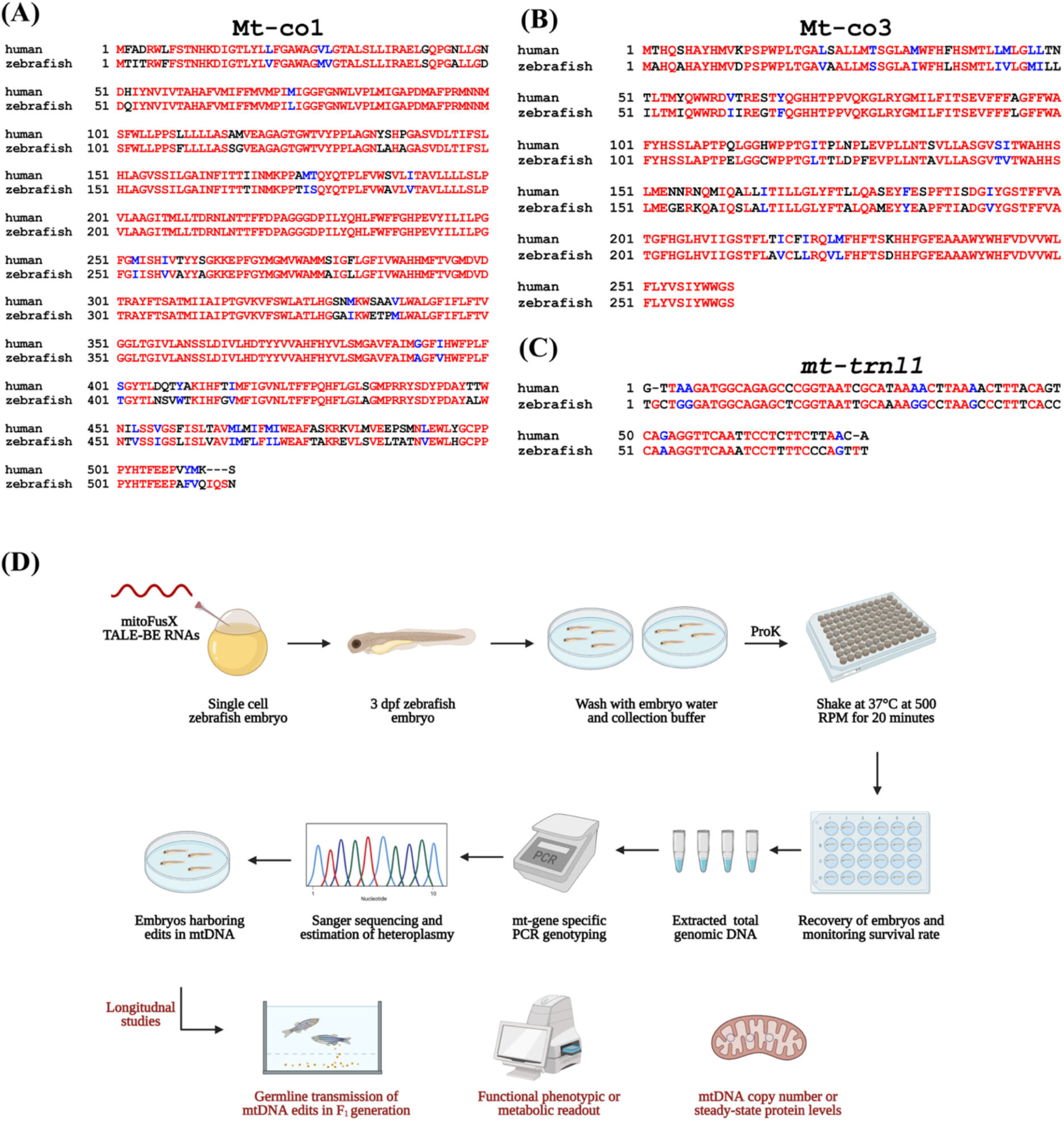
Schematic to prioritize base editing targets and non-invasively genotype mtDNA from zebrafish embryos. **(A and B)** Targets identified in the mitochondrial protein-coding genes, *mt-co1* and *mt-co3*, were prioritized to test the editing efficiency of FusXTBE *in vivo*. Amino acid alignments are shown for the human MT-CO3 and MT-CO1 proteins with zebrafish Mt-co1 and Mt-co3 proteins. **(C)** The mitochondrial-encoded tRNA gene, *mt-trnl1*, was prioritized to introduce base edits. Sequence alignment of human MT-TRNL1 and zebrafish orthologue *mt-trnl1* are shown. Red indicates nucleic acid sequences fully conserved between the two species. **(D)** Longitudinal study design assay for potential functional phenotype of the engineered edits in the zebrafish mtDNA locus. Zebrafish embryos are gently shaken in the genotyping buffer supplemented with proteinase K. Following recovery of the embryos, DNA is isolated from the shed cells, and mitochondrial genotyping is conducted for the target locus. Positive embryos can then be screened for germline transmission or different functional and molecular studies. Figure 4b was created using Biorender.com. For figure 4A, sequences were aligned using performed by T-COFFEE multiple sequence alignment webserver and color-coded using BoxShade server.

*mt-co1* BE RNAs was designed to edit the cytosine (C_8_) on the reverse strand at the m.C7106 of zebrafish mtDNA (Figure 5A). m.G7106A edit on the forward strand or (C-to-T transition on the reverse strand) leads to the loss of a restriction site for the enzyme BstYI. Amplification of the target locus from the extracted DNA followed by mitochondrial genotyping revealed the presence of undigested band (431 bp) in the case of injected embryos as compared to the digested products (265 and 166 bp observed in the non-injected control embryos) (Figure 5B). Sanger sequencing files (.ab1) were then assessed using EditR software^21^ to quantify the level of editing injected embryos. This analysis demonstrated that FusXTBE was highly efficient *in vivo* as compared to *in vitro*, showing editing efficiencies as high as up to 90% (Figure 5C). More than 70% of the injected embryos exhibited mutant mtDNA heteroplasmy. Analysis of the protospacer region revealed that the editing happened in both strands in the pyrimidine stretch present in the protospacer region, with efficiency in the range of 60-80% (Figure 5C-D) in individual zebrafish embryos.

**Figure 5:**
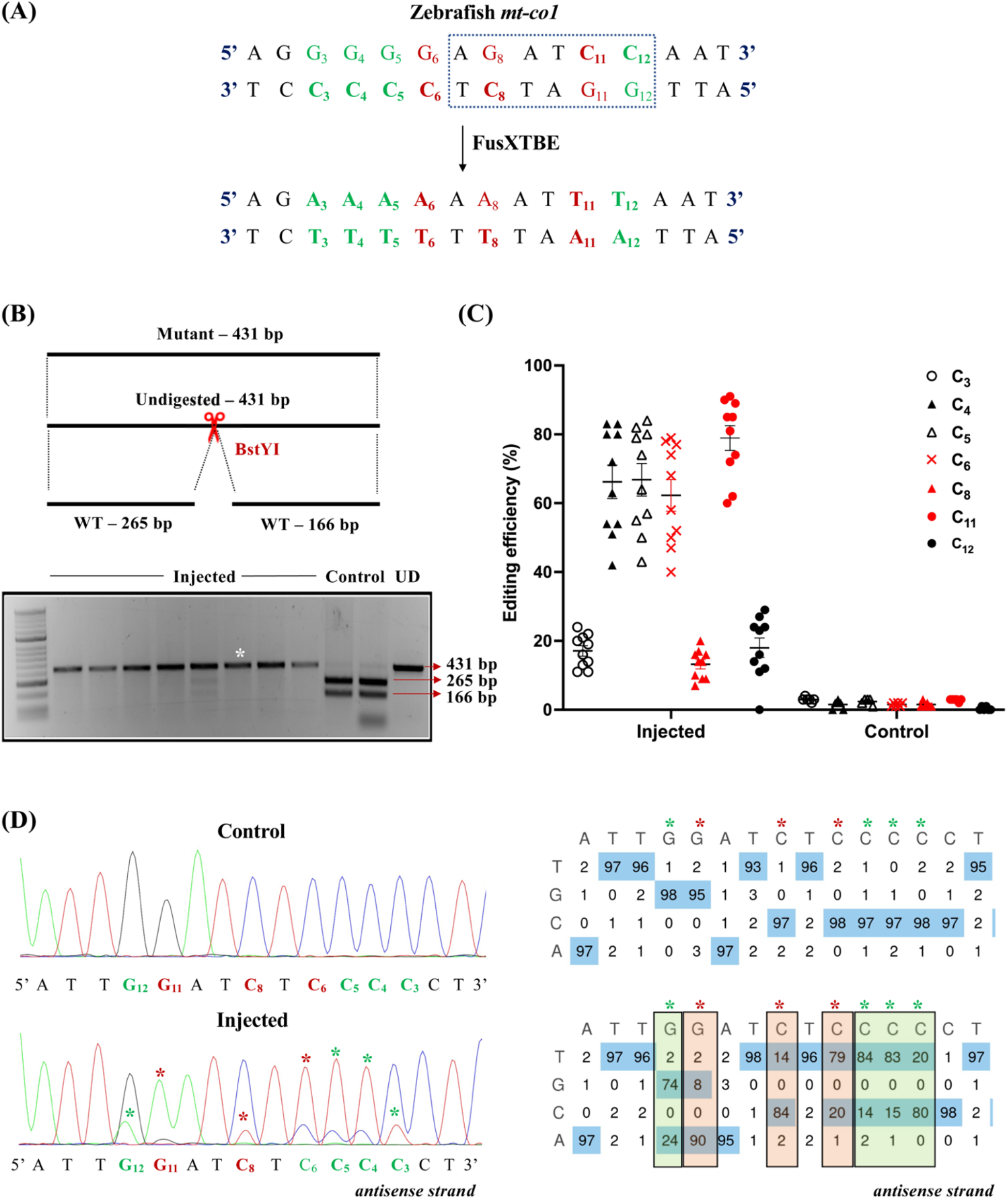
Introducing base edits in the zebrafish *mt-co1* gene using the FusXTBE system. **(A)** Protospacer region in the mt-co1 locus highlighting the potential editing sites. Restriction site for the enzyme BstY1 is marked by dotted rectangle. **(B)** Screening of embryos harboring mutations by RFLP. C-to-T edits in the protospacer region leads to loss of recognition site for BstyI restriction enzyme. PCR amplicon from non-injected control embryos show the presence of two wild-type digested products. PCR amplicons from the injected embryos indicate the presence of an undigested band marked by white asterisk (*). Each lane represents an individual embryo. **(C)** Editing efficiency for the favorable 5’TC sites present in the protospacer region both in injected and non-injected control zebrafish embryos. Specific and non-specific target nucleotides are indicated by red and black colored circles, respectively. Each data point represents an individual zebrafish embryo. Data is represented from independent experiments. **(D)** Representative chromatogram of the control and injected embryos. Asterisk (*) denotes the site of edit with corresponding editing percentage (C-to-T or G-to-A). Specific and non-specific edits are denoted by red and green asterisks (*), respectively. Chromatograms and editing table plot were obtained using EditR.

To leverage the potential of this construct in studying loss of function studies for mitochondrial genes, we chose to test the system against the site m.C10215 (C_10_) present in the *mt-co3* gene (Figure 6A). A C-to-T edit at this position creates a new premature termination codon, UAA, leading to a truncated protein of 160 amino acids [p.(Gln161*)], as determined by *in silico* analyses. Embryos were scored based on the gain of restriction site for the enzyme MseI. RFLP analyses from the injected animals revealed the presence of digested products at the expected size of 159 and 152 bp along with the 311 bp undigested WT band (Figure 6B). RFLP analysis was confirmed by Sanger sequencing, wherein RFLP positive embryos displayed 85% mutant heteroplasmy (Figure 6C-D). We did not observe any non-specific C-to-T edits in the protospacer region (C_14_ and C_16_).

**Figure 6:**
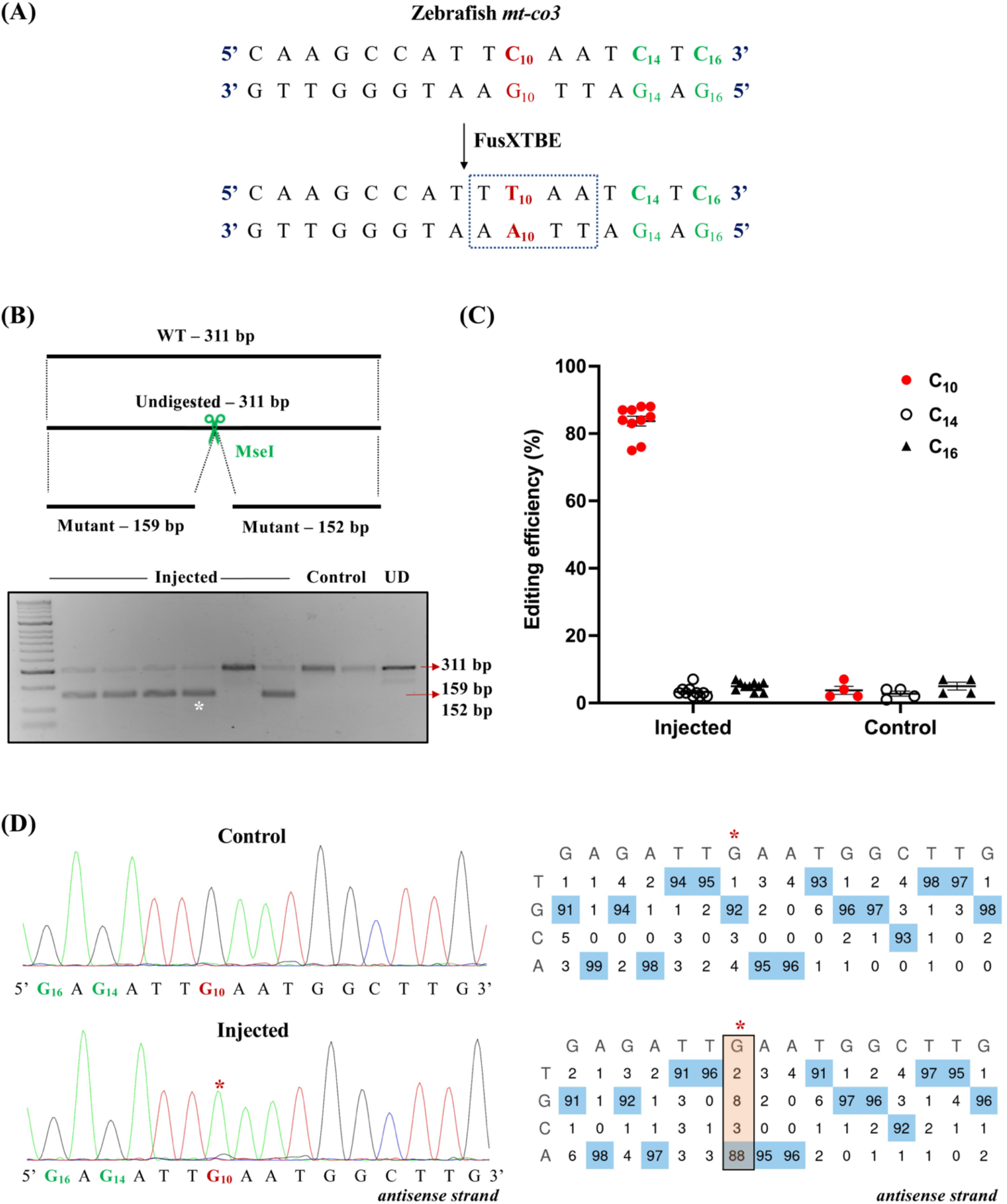
Engineering precise base edits in vivo in the zebrafish *mt-co3* gene. **(A)** Sequence of the potential editing sites in the *mt-co3* protospacer region. The resulting gain of the restriction site for the enzyme MseI post editing event is marked by a dotted rectangle. **(B)** Screening of embryos harboring mutations by RFLP. C-to-T edits in the protospacer region leads to gain of recognition site for the MseI restriction enzyme. PCR amplicon from non-injected control embryos show the presence of uncut product (311 bp). PCR amplicons from the injected embryos indicate the presence of expected 159 and 152 bp digested bands marked by a white asterisk (*). Each lane represents an individual embryo. **(C)** Editing efficiency for the target C_10_ present in the protospacer region both in injected and non-injected control zebrafish embryos. Specific (C_10_) and other DddA_tox_ amenable (C_10_ and C_16_) nucleotides are indicated by red and black colored circles, respectively. Each data point represents an individual zebrafish embryo. Data is represented from independent experiments. **(D)** Representative chromatogram of the non-injected (control) and injected embryos. Asterisk (*) denotes the site of edit with corresponding editing percentage (C-to-T or G-to-A). Chromatograms and editing table plot were obtained using EditR.

To confirm that our results were not specific to just protein-coding genes, we decided to test our system against the zebrafish mitochondrial tRNA gene, *mt-trnl1* (Figure 7A). We chose a site that was conserved across the human and zebrafish genomes. A C-to-T transition at the target site m.G3739 (C_5_ on the reverse strand) created a loss of restriction site for the enzyme SacI (Figure 7B). Individual embryos that displayed the loss of restriction site corresponding to the uncut band at 363 bp (Figure 7B) harbored 60-70% mtDNA mutant heteroplasmy levels for the target locus (Figure 7C-D). We also observed another edit at m.C3744 (C_10_) in the protospacer region with comparable mutant heteroplasmy. This observation cannot be deemed as off-target since any C which is preceded by T at the 5’ end is observed to be amenable to base editing by the DdCBE system as reported previously^17^.

**Figure 7:**
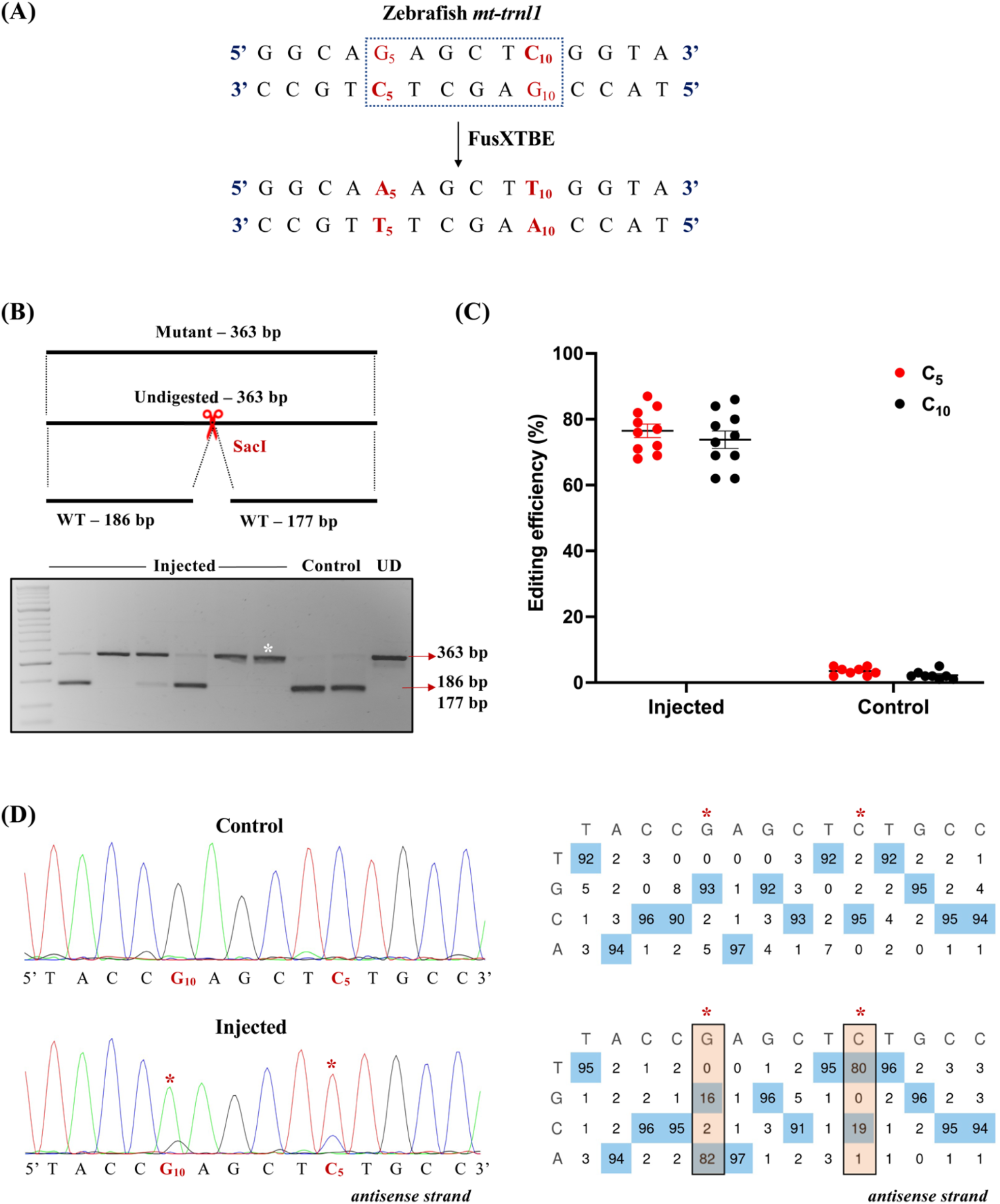
Modeling mutations in the zebrafish *mt-trnl1* gene. **(A)** Protospacer region highlighting the putative cytosine targets in the *mt-trnl1* locus. Restriction site for the enzyme SacI is marked by a dotted rectangle. **(B)** Screening of embryos harboring mutations by RFLP. C-to-T editing event in the target locus leads to loss of restriction site for the SacI restriction enzyme. Presence of undigested WT bands of expected size, 186 and 177 bp (marked by red arrow), is observed in control embryos. PCR amplicons from the injected embryos show the undigested band, indicated by the loss of restriction site post-editing (marked by white asterisk*). Each lane represents an individual embryo. **(C)** Editing efficiency for the favorable 5’TC sites present in the protospacer region both in injected and non-injected control zebrafish embryos. Specific and nonspecific target nucleotides are indicated by red and black color circles, respectively. Each data point represents an individual zebrafish embryo. Data is represented from independent experiments. **(D)** Representative chromatogram of the control and injected embryos indicating the percentage of editing at each cytosine residue in the protospacer region. Asterisk (*) denotes the site of edit with corresponding editing percentage (C-to-T or G-to-A). Specific and non-specific edits are denoted by red and green asterisks (*), respectively. Chromatograms and editing table plot were obtained using EditR.

## Discussion

Mitochondrial DNA engineering had witnessed significant advancements in the past decade due to the advent of mitochondrially targeted nucleases that leveraged the potential of the absence of double stranded break repair in shifting the heteroplasmy from mutant to wild type mtDNA levels^11, 12, 15^. However, the utility of this toolkit could not address treating mutant mtDNA homoplasmy or introducing precise edits in the mitochondrial genome. Mok *et al*., recently described a bacterial toxin, cytosine deaminase that catalyzes the conversion of C•G to T•A in the mtDNA^17^. This obligate two-component system utilizes two separate non-toxic halves of DddA_tox_ fused to TALE monomers to introduce single nucleotide changes in the mtDNA *in vitro*. This system was not readily accessible due to the nature of the TALE construction approach used and did not include any corresponding informatics-based genomic analyses for exploring the potential gene editing outcomes. We built upon this initial architecture and extended the application of mtDNA base editing to model different mutations in mitochondrial genes implicated with clinical manifestations and to leverage it as a tool to understand basic mitochondrial biology.

As described previously, investigating a single target would require a user to test four different combinations of the DdCBE system (left-G1397-N and right-G1397C, left-G1397-C and right-G1397N, left-G1333-N and right-G1333C or left-G1333-C and right-G1333N)^17^. This system does open an avenue to model or treat single nucleotide pathogenic variations in animal and cellular models. However, traditional methods of TALE assembly either by conventional Golden Gate cloning or via expensive gene blocks impede the progress that the community can make in establishing mtDNA disease animal models. To circumvent these challenges, we integrated our rapid one-step Golden Gate FusX TALE assembly system for use with the DdCBE base editor. We created the mitochondrial FusX TALE base editor (FusXTBE) architecture, where split halves of DddA_tox_ and UGI were fused with a modular FusX^19^ compatible with MTS-TALE module. The FusX system consists of 340 plasmids, which are pre-assembled trimers (spanning across six libraries), to generate 14.5- to 16.5-mer TALE repeats in a single Golden Gate reaction with a 3-days turnaround time. This toolbox offers an additional advantage of being compatible with mitochondrially and nuclearly targeted TALE scaffolds. The FusX system can be automated and integrated to a high throughput robotic liquid handling device, thereby scaling up the number of TALE base editors that can be assembled with minimal human error. This system is available at Addgene and has been successfully utilized for nuclear and mitochondrial gene editing applications, both *in vitro* and *in vivo*, with high efficiency^16, 19^.

The DdCBE system is known to edit C in a 5’T**C** sequence specific manner on any of the strands. To aid in interrogating different potential base editing sites, we developed TALE Writer, a customizable Python-based script. TALE Writer not only allowed predicting 5’T**C** sites across the human and zebrafish mitochondrial genomes. It also provided the top-ranked combinations of TALE arms and spacer regions specific for each identified site. While we identified more than 1,500 such potential 5’T**C** sites in each of the two species, we were further interested in modeling loss of function studies for mitochondrially encoded protein-coding genes. Loss of function or translational defects due to the creation of premature stop codons in a protein-coding gene will allow us to elucidate its role in mitochondrial disease biology. TALE Writer identified more than 85 such sites in each of the interrogated genomes and can be used to model clinically significant loss of function mtDNA variations in the protein-coding genes, both in cellular and animal models (such as zebrafish). The computation time of the script is usually negligible (the output is virtually immediate) from the moment a user provides an input sequence for a protein-coding gene to obtaining an output with the top-ranked FusX compatible TALE sequences. All the targets used in this study were identified by the algorithm and were experimentally validated in different *in vitro* and *in vivo* model systems.

As a quantitative test of the FusX TALE modular cassette to DddA_tox_ as a base editor, we compared our system with the previously published DdCBE system for the same mtDNA locus^17^. It was notable from our findings that FusXTBE demonstrated no significant difference in activity in this *in vitro* assay (Figure 3B). This served as a reference test system for expanding applications including additional work *in vivo*. For example, the modular linker present in FusXTBE between the RVD domain and DddA_tox_ enables changes in the potential editing window in the protospacer region. FusXTBE also offers an opportunity to drive heteroplasmy levels well beyond 30% for other mtDNA targets in mammalian cell lines.

Multi-systemic clinical manifestations and varying mtDNA heteroplasmy from tissue to tissue in the case of mitochondrial disorders cannot be replicated by cells in a dish. Therefore, we explored the opportunity of testing the FusXTBE system *in vivo*. Zebrafish is a suitable animal model for studying mitochondrial disorders due to the 65% sequence conservation with human mtDNA as well as the shared mitochondrial-specific genetic code, strand-specific nucleotide frequencies and fully syntenic gene order^25, 26^. Microinjection of FusXTBE RNAs in single cell zebrafish embryos not only enhances the cellular delivery but gives a rapid turnaround time to genotype positive embryos within 3 days post fertilization (3 dpf). The poor correlation of phenotype to genotype and stochastic mode of transmission makes it challenging to study the biology of the mtDNA-encoded disorders. To overcome this technical hurdle, we adopted a non-invasive enzymatic genotyping approach^22^ to screen for embryos harboring positive edits. This approach allowed us to subject the animals to functional assessment or monitoring the germline transmission of the edits in the F1 generation. This assay involves the extraction of DNA from the cells that are shed after an event of modest shaking, ensuring optimal survival. This assay was originally described earlier for the nuclear genotyping^22^ and with some modifications, we have adapted this assay to genotype mtDNA from zebrafish embryos. High survival rate of injected embryos, i.e., 80%, and successful amplification of the mtDNA RFLP, coupled with rapid TALE assembly, accelerates the process to model various pathogenic mutations *in vivo*.

One of the rationales to prioritize targets in zebrafish was to look for genes involved in clinically associated symptoms and those that are indispensable for mitochondrial function. Mutations in *MT-CO1* and *MT-CO3* are associated with conditions such as Leber optic atrophy, cytochrome c oxidase deficiency^27, 28^. MELAS disease is primarily associated with mutations in the gene *MT-TRNL1* and is presented with symptoms such as mitochondrial encephalomyopathy, lactic acidosis, and stroke-like episodes^4^. Initially, we chose a pyrimidine rich target on the zebrafish *mt-co1* gene to identify the editing window within the protospacer region. Genotyping analyses revealed that approximately 70% of the injected embryos displayed editing frequencies of up to 90%. Interestingly, we also observed several non-specific C-to-T edits having a 5’T**C**C sequence context, which may be due to the accessibility of the DddA_tox_ molecule to the unstacked pyrimidine strand during the editing event.

Gaining insights from these observations, we chose a target in the *mt-co3* locus to introduce a premature termination codon upon a successful C-to-T edit at the m.C10215 position, thereby truncating the amino acid sequence, instead of the usual glutamine at that location. Sequence specificity for such targets is of utmost importance to avoid any non-specific amino acid substitutions either upstream or downstream of the target nucleotide. Consistent with the high editing efficiency percentage observed for the previous target, frequencies of C•G to T•A transitions ranged between 75-88% for the injected individual embryos. Such high percentages of precise edits *in vivo* generated with FusXTBE in zebrafish mtDNA allow us to understand the functional consequence of the truncation of mitochondrial Co-3 protein in the context of mitochondrial biology. None of these mutations were embryonically lethal, allowing us to monitor the germline transmission of these edits in subsequent F1 generations during the future course of the study. It will be interesting to see if such high percentage of edits are able to withstand the oocyte selection of mutant mtDNA as observed previously with low heteroplasmy of mtDNA large-scale deletion in zebrafish^16^. Recently, a study describing mice editing with a custom designed DdCBE demonstrated base edits in mice F0 and F1 embryos albeit with maximum mtDNA editing efficiency up to 57% in F0 generation for one of the loci. Two pups born to the mutant F0 female mouse harboring edits exhibited edits at a frequency that ranged between 6-26%^18^.

Our analysis documented C•G to T•A transitions in another target, *mt-trnl1*, ortholog of human gene *MT-TRNL1* that is implicated in the pathogenicity of the MELAS. Approximately 70% of MELAS cases are attributed to the m.A3243G^29^ mutation in the human *MT-TRNL1* gene. However, modeling this exact variation was not possible utilizing the DddA_tox_ system due to the lack of a 5’T**C** in the target locus. We adopted a sequence similarity and secondary structure alignment informatic analyses to prioritize targets in the zebrafish locus (data not shown). One of the targets identified in the zebrafish *mt-trnl1* gene was upstream to the conserved nucleotide to m.A3243 of the human orthologue. In 70% of the injected embryos, we observed C-to-T edits at m.G3744 (C_10_) in the range between 69-82%. We did notice another edit with similar percentage at m.G3739 (C_5_) within the protospacer window. This non-specific edit could be masked by using our FusX Extender series to design 17-mer TALE repeats that could bind to this nucleotide and make it inaccessible for cytidine deaminase activity. This improved FusXTBE architecture could aid in eliminating non-specific edits near the targeted cytosine residue.

In summary, our study reports the first incidence of *in vivo* editing of mtDNA in zebrafish using a programmable FusX compatible TALE base editor. Rapid one-tube FusXTBE system with TALE Writer enables the facile design of TALE targets against potential base editing (5’TC) sites in human and mitochondrial genomes. The FusXTBE system demonstrated more than 85% C•G to T•A transitions in individual zebrafish embryos for both protein-coding and tRNA genes. Studies measuring the germline transmission of high mutant mtDNA heteroplasmy load and genotype to phenotype correlation will shed more light on the underlying mitochondrial disease biology of mtDNA variations.

## Conclusion

Mitochondrial medicine was galvanized by the recent discovery of the DdCBE system enabling single nucleotide edits in the mitochondrial genome. In this study, we build upon this paradigm to extend the base editing *in vivo* using the rapid one-tube Golden Gate TALE assembly system, FusXTBE. Our programmable system also provides an adaptable algorithm to predict potential base editing sites in the human and zebrafish genome based on known editing guidelines, facilitating the design of TALE targets, and accelerating turnaround time. The FusXTBE system has demonstrated editing efficiencies of more than >85% in the mtDNA locus *in vivo*, offering unique insights into using this automatable and high-throughput toolbox to model or correct pathogenic mutations in the mitochondrial genome in cellular and animal models. The findings presented in this study opens an avenue to explore the biology of the mitochondrial genome and its implications in mitochondrial disorders.

## Acknowledgment

The authors thank Dr. David Liu, Harvard University for kindly providing the DdCBE plasmids through Addgene (Kit #1000000063). Authors acknowledge Dr. Eric Schon, Columbia University for providing valuable input during the preparation of this manuscript. We would also like to thank Kavini Nanayakkara and the Mayo Clinic Zebrafish Facility staff.

## Author contributions

The idea was conceived by KJC and SCE. Manuscript was written by AS, SRC and SCE. Experiments were executed by AS, BK, SRH, BLK and SRC with experimental guidance from KJC and SCE. Data analysis was completed by AS, BK and SRC. FusX Extender series was assembled by ZWJ, RPC and KJC. Manuscript was reviewed and edited by AS, BK, SRC, SRH and SCE. Authors declare no competing interests.

## Data availability

Plasmids are available on request. FusX assembly kit is available through Addgene.

## Supplementary table

**Supplementary table 1:**
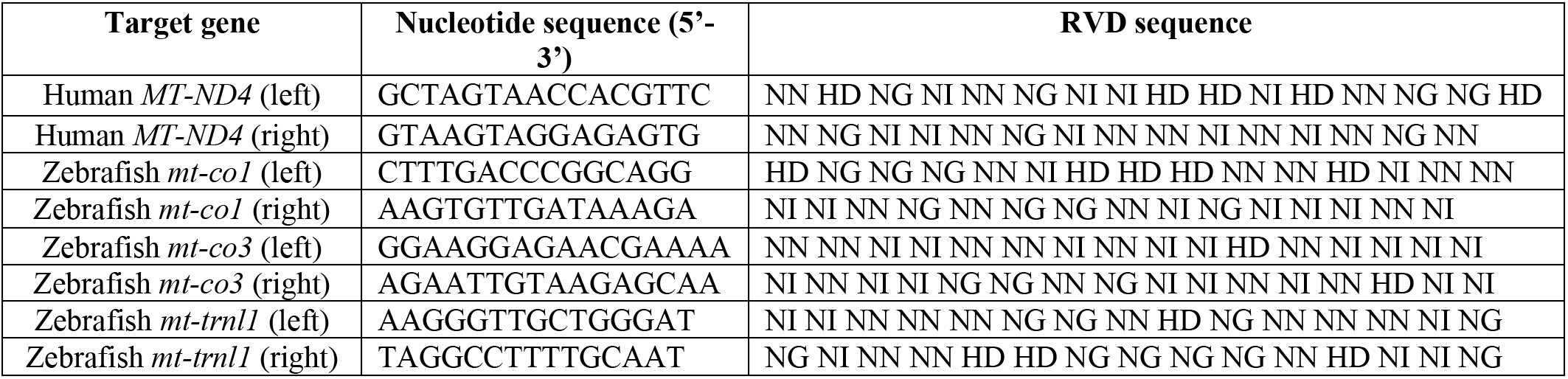
RVD sequence for mtDNA loci targeted by FusXTBE.

**Supplementary table 2:**
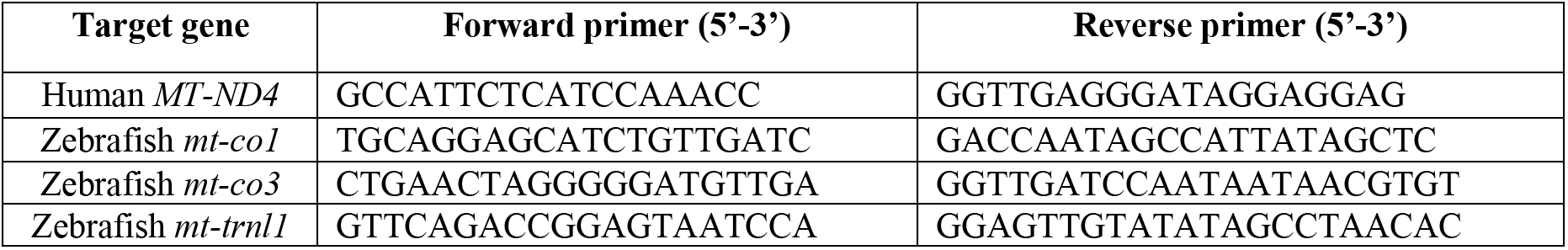
List of oligonucleotides used in the study.

## Supplementary figure

**Supplementary figure 1:**
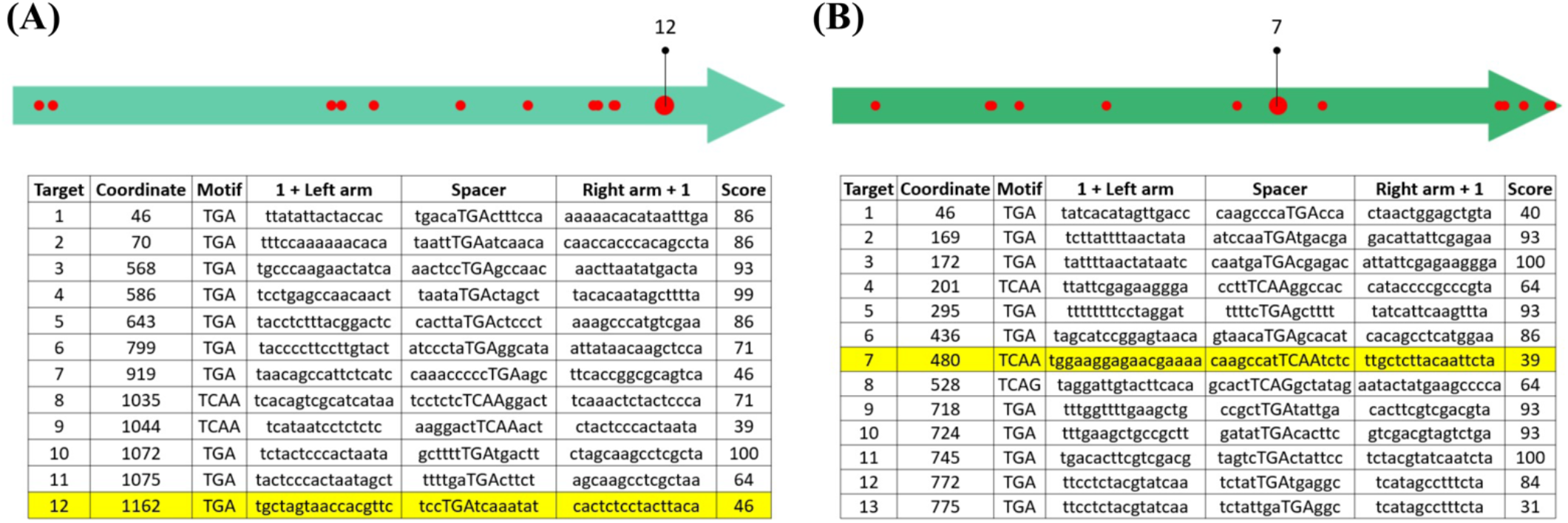
TALE Writer predicted targetable sites for PTC induction in the (A) human gene *MT-ND4* and the (B) zebrafish gene *mt-co3*. Horizontal arrows represent the open reading frames for each gene. The tables shown below correspond to raw output generated by TALE Writer. For each target site, the following information is given: (I) The coordinate of the target with respect to the start site of the input sequence, (II) the target motif (i.e., TGA, TCAA, or TCAG), (III) the target sequence of the left TALE (always preceded by an extra 5’T0), (IV) the spacer sequence (with only the target motif in capital letters), (V) the sequence complementary to the target sequence of the right TALE (plus an additional 3’A, which corresponds to a 5’T0 in the actual TALE target sequence), and (VI) a score with a minimum value of 31 and a maximum value of 100. TALE Writer generates the top-ranked design permutations for each target, but other design permutations are stored and accessible. (**A)** Utilizing initial selected design parameters, TALE Writer identified a total of 12 target sites amenable to PTC induction in the human mitochondrial gene *MT-ND4*. In the light green arrows, each predicted PTC site is represented by a red circle located at its corresponding local coordinate. Site number 12 (big red circle on the arrow, highlighted row in the table) was selected for experimental validation. **(B)** Based on the same design parameters, TALE Writer identified a total of 13 target sites amenable to PTC induction in the zebrafish mitochondrial gene *mt-co3* (red circles). Site number seven (big red circle on the arrow, highlighted row in the table) was selected for experimental validation.

